# Novel multi-omics deconfounding variational autoencoders can obtain meaningful disease subtyping

**DOI:** 10.1101/2024.02.05.578873

**Authors:** Zuqi Li, Sonja Katz, Edoardo Saccenti, David W. Fardo, Peter Claes, Vitor A.P. Martins dos Santos, Kristel Van Steen, Gennady V. Roshchupkin

**Author notes:** Authors contributed equally (Zuqi Li, Sonja Katz).

## Abstract

Unsupervised learning, particularly clustering, plays a pivotal role in disease subtyping and patient stratification, especially with the abundance of large-scale multi-omics data. Deep learning models, such as variational autoencoders (VAEs), can enhance clustering algorithms by leveraging inter-individual heterogeneity. However, the impact of confounders - external factors unrelated to the condition, e.g. batch effect or age - on clustering is often overlooked, introducing bias and spurious biological conclusions. In this work, we introduce four novel VAE-based deconfounding frameworks tailored for clustering multi-omics data. These frameworks effectively mitigate confounding effects while preserving genuine biological patterns. The deconfounding strategies employed include: i) removal of latent features correlated with confounders ii) a conditional variational autoencoder, iii) adversarial training, and iv) adding a regularization term to the loss function. Using real-life multi-omics data from TCGA, we simulated various confounding effects (linear, non-linear, categorical, mixed) and assessed model performance across 50 repetitions based on reconstruction error, clustering stability, and deconfounding efficacy. Our results demonstrate that our novel models, particularly the conditional multi-omics VAE (cXVAE), successfully handle simulated confounding effects and recover biologically-driven clustering structures. cXVAE accurately identifies patient labels and unveils meaningful pathological associations among cancer types, validating deconfounded representations. Furthermore, our study suggests that some of the proposed strategies, such as adversarial training, prove insufficient in confounder removal. In summary, our study contributes by proposing innovative frameworks for simultaneous multi-omics data integration, dimensionality reduction, and deconfounding in clustering. Benchmarking on open-access data offers guidance to end-users, facilitating meaningful patient stratification for optimized precision medicine.

## 1 Introduction

Unsupervised learning, in particular clustering, focuses on subgrouping individuals based on their intrinsic data structures, therefore playing an essential role in tasks like disease subtyping and patient stratification. In the realm of biology and medicine, where large-scale multi-omics data, including genomics, transcriptomics, and epigenomics, is prevalent, deep learning models can enhance clustering algorithms. Their ability to reduce the dimensionality of complex data allows clustering algorithms to more effectively explore the heterogeneity between patients. Underscoring the utility of deep learning models, in particular autoencoders, in terms of data integration, dimensionality reduction, and handling a multitude of heterogeneous input data, Simidjievski et al. recently benchmarked various variational autoencoder models for multi-omics data [24].

Although patient stratification with deep learning methods are gaining traction in genomic data applications, they are often susceptible to external influences that are unrelated to the condition of interest. One severe limitation is the entanglement of biologically meaningful signals with variables unrelated to the inherent structure that one is interested in, i.e. technical artifacts, random noise from measurements, or other biological factors such as sex, ethnicity, and age (Figure 1a). These factors, referred to as confounders in the context of unsupervised learning, may cause clustering algorithms to form subgroups based on irrelevant signals, which may ultimately lead to spurious biological conclusions [5, 11].

**Figure 1.**
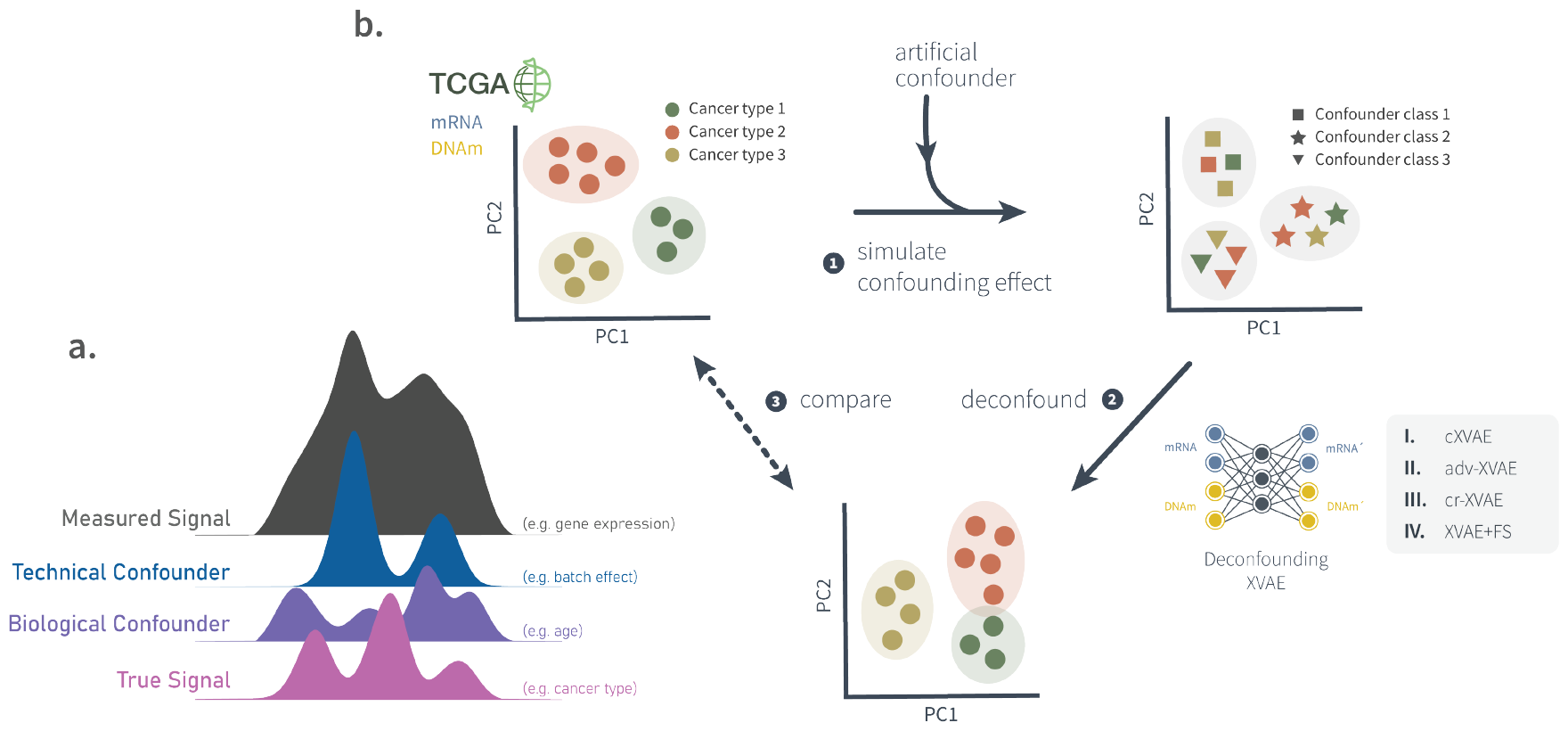
**a**. A simplified graphical representation of a measured signal (gray) which is a mix of independent sources such as the true signal (pink), a biological confounder (purple), and a technical confounder (blue). Note the difficulty of extracting the true signal from the measured additive signals. **b**. Graphical summary of the work conducted in this study. (1) Based on multi-omics pan-cancer TCGA data (section 2.1) different confounding effects were simulated (section 2.2). (2) Subsequently, four different deconfounding VAE frameworks (section 2.4) were trained on the the artificially confounded data. (3) The obtained deconfounded data was compared to the original, un-confounded input data in terms of clustering stability and deconfounding capabilities (section 2.6).

Conventional strategies to account for and mitigate confounders involve training linear regression per feature against the confounder and take the residual part during preprocessing [22] or adjustments like pruning predictive dimensions after model training [23]. Conditional variational autoencoders (cVAE) have been used to create normative models considering confounding variables, such as age, for neurological disorders [17]. Dincer et al. proposed adversarial training to derive expression embeddings devoid of confounding effects [7], expanded upon by the single-cell Generative Adversarial Network (scGAN) for batch effect removal [2]. Liu et al. used a regularization term in the autoencoder’s loss function to minimize correlation between latent embeddings and confounding bias [18]. Despite their methodological diversity, these methods have only been validated to work effectively on data from a single omics source and are not tailored towards disease subtyping and patient stratification.

To address this gap, we propose four novel VAE-based deconfounding frameworks for clustering of multi-omics data, utilising the i) removal of latent features correlated with confounders ii) a conditional variational autoencoder [17] iii) adversarial training [2, 7], and iv) adding a regularisation term to the loss function [18] as deconfounding strategies. To objectively assess whether our models can remove out-of-interest signals and find a patient clustering unbiased by confounding signals, we pplied and evaluated our models on gene expression and DNA methylation pan-cancer data from The Cancer Genome Atlas (TCGA) program which we augmented with artificial confounding effects. In total, we simulated four different types of confounders, including linear, non-linear, categorical, and a mixture thereof, resembling realistic confounders such as age (linar, non-linear) [6, 15, 26], BMI (non-linear) [20], or batch effects (categorical) [7, 11].

The contribution of our study is as follows:

- Four novel multi-omics clustering models based on VAE and different deconfounding strategies are presented.
- We highlight that various deconfounding techniques address confounded clustering in distinct ways, often overlooked within the algorithm’s framework.
- Different confounding effects are simulated on the real-life TCGA dataset to demonstrate the influence of confounders on clustering and underscore the necessity for deconfounding models.
- Readers are provided with guidelines detailing strengths and limitations of each approach, along with suggestions on selecting an appropriate framework fitting their purposes.

## 2 Materials and methods

### 2.1 Data collection & preprocessing

This study utilized data collected within The Cancer Genome Atlas project (TCGA) [29]. Gene expressions (mRNA) of 4333 patients and DNA methylations (DNAm) of 2940 patients from six different cancer types, including BRCA, THCA, BLCA, LUSC, HNSC, and KIRC, were downloaded using the R package TCGAbiolinks [4]. The subsequent filtering step removed patients with (i) only a single data type available, (ii) missing clinical metadata, (iii) “american indian” or “alaska native” ancestry, and (iv) unknown tumor stage, resulting in a total of 2547 patients. The preprocessing of mRNA and DNAm data included the removal of probes (i) not shared across all cancer types, (ii) with missing values, and (iii) with 0 variance across all included patients, resulting in 58456 mRNA and 232088 DNAm features. To reduce the number of input features, we only considered the 2000 probes showing the largest variance across patients for each data type, resulting in a final data set of 2547 patients and 4000 features. This reduction strikes a balance between the number of features included and biological variability addressed and is in line with other clustering works on TCGA data [3]

### 2.2 Simulation of confounders

To imitate common confounding scenarios in real-life clustering applications we simulated linear, squared, categorical confounders, and a mixture thereof, resembling e.g. ageing [6, 15, 26], BMI [20], or batch effects [7, 11]. These confounders hinder the true or biologically meaningful clustering by intrinsically affecting the data structure in an unwanted way and possibly leading to a confounded clustering.

Here we denote the mRNA data as *X*_1_ ∈ ℝ ^*n×p*^ and DNAm data as *X*_2_ ∈ ℝ^*n×q*^, where *n*,*p, q* are the number of patients, gene expressions, and DNA methylations, respectively. We first rescaled every mRNA and DNAm feature to the range [0, 1] to avoid large ratios between the raw feature and the confounding effect. A visualisation of all confounding effects can be found in the Supplementary Methods.

#### 2.2.1 Linear confounder

We uniformly generated a random numeric confounder **c** ∈ ℝ ^*n*^ with discrete values {0, 1, 2, 3, 4, 5}, leading to a confounder clustering of six classes. Its linear effect on each individual is **c** + 5 and a random weight for each feature was multiplied with it:

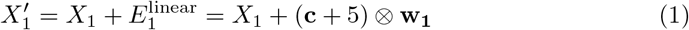

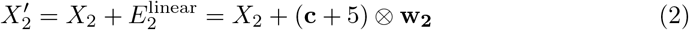

, where ⊗ denotes the outer product between two vectors, and **w**_**1**_ *∈* ℝ^*p*^ ∼ *U* (0, 0.1), **w**_**2**_ ∈ ℝ^*q*^ ∼ *U* (0, 0.2). We chose the uniform distribution of **w**_**1**_ to range from 0 to 0.1 so that the total linear effect would range from 0 to 1, having the same scale as *X*_1_. We increased the upper bound of **w**_**2**_ to 0.2 due to our observation that *X*_2_ is less sensitive to linear confounders.

#### 2.2.2 Non-linear confounder

Non-linear effects were simulated in a similar way to linear effects. However, to mimic a non-linear confounder, as observed in, e.g. the significant quadratic association between body mass index and colon cancer risk [20], we considered adding an element-wise squared confounding effect **c**^2^ on the features:

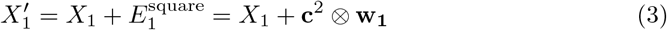

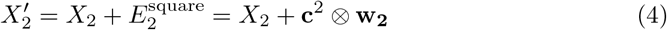

, where **w**_**1**_ ∼ *U* (0, 0.04), **w**_**2**_ ∼ *U* (0, 0.04). The distribution of **w**_**1**_ and **w**_**2**_ was also determined based on the scale of *X*_1_ and *X*_2_.

#### 2.2.3 Categorical confounder

A categorical confounding effect was achieved by shifting patients with the same confounder class to a distinctive direction in the feature space. More specifically, we first sampled six shifting vectors from *U* (0, 1) corresponding to six different confounder classes, and patients were randomly assigned to each of the six categories. As a result, *C*_1_ ∈ ℝ^*n×p*^ denotes the concatenation of shifting vectors of every patient for gene expression, while *C*_2_ ∈ ℝ^*n×q*^ for DNA methylation, both are matrices. The categorical confounder is therefore the membership of all individuals in the six classes. A typical example of categorical confounders for clustering could be batch effects caused by collecting data from different centers [7, 11]. Then the confounded features were created via:

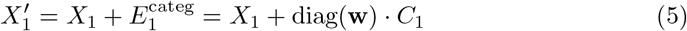

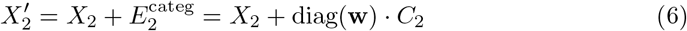

, where diag(*·*) converts a vector into its corresponding diagonal matrix. Different from the case of a numeric confounder, the weight vector **w** ∈ ℝ^*n*^ ∼ *U* (0, 1) of the categorical confounder indicates to what extent every patient was shifted so that patients would have various strength of association with their confounded class.

#### 2.2.4 Mixed confounder types

Real-life data analyses are likely affected by multiple confounders of different kinds, for instance, many cancer studies correct for age, age squared, education, etc. jointly in their models [15, 26]. Here we simulated a mixed confounding effect of linear, non-linear, and categorical confounders as described below:

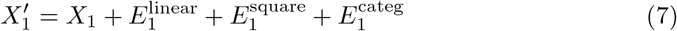

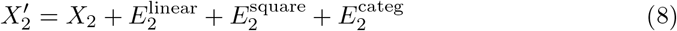

, where 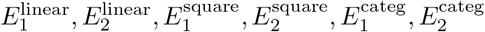 represent the second term in Formula (1-6), respectively.

### 2.3 Variational autoencoder for data integration (XVAE)

A variety of different VAE architectures exist for the purpose of data integration, as extensively compared by Simidjievski et al. [24]. In this study, we utilize one architecture recommended by the respective authors, namely the X-shaped Variational Autoencoder (XVAE) (Figure 2). This architecture merges the heterogeneous input data sources into a combined latent representation *z* by learning to reconstruct each source individually from the common representation. Here we consider only two data types of a single datapoint *x*_1_ and *x*_2_, and the loss function of XVAE is as follow:

**Figure 2.**
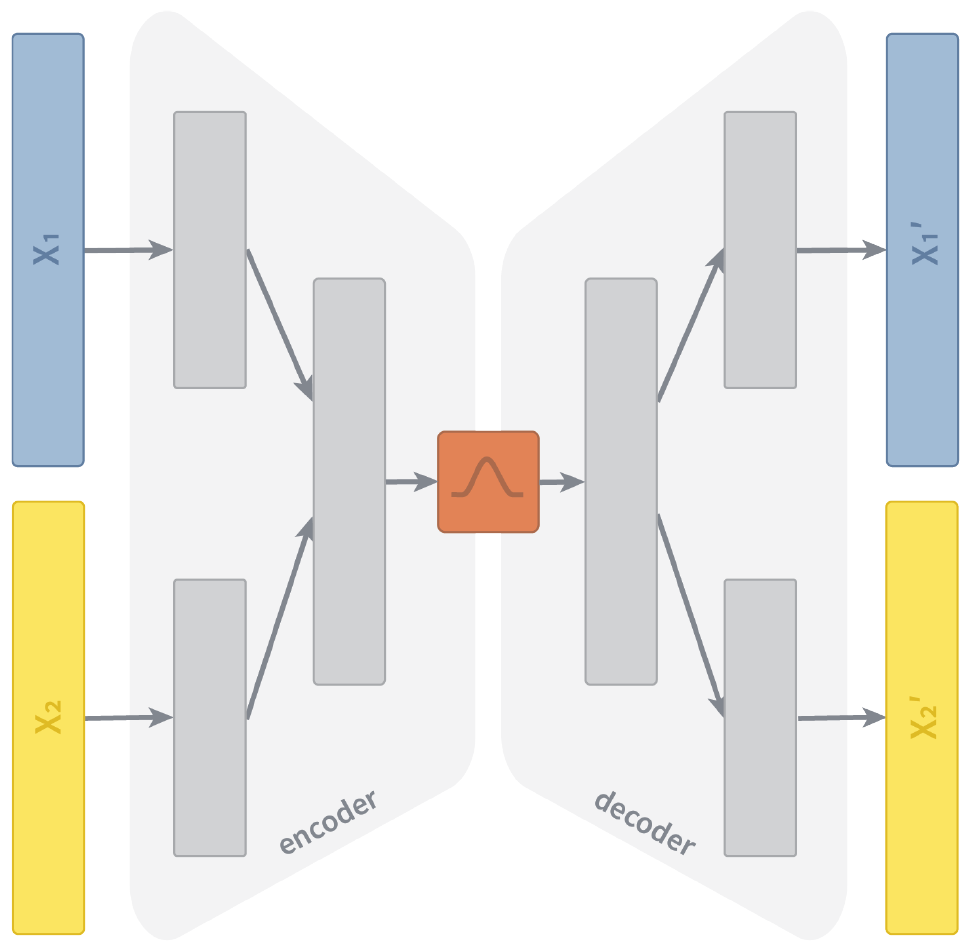
Schematic representation of an X-shaped Variational Autoencoder (XVAE). The two input layers (*X*_1_, *X*_2_) denote the two omics dimension used in this study, namely gene expression and DNA methylation. The encoder consists of contiguous hidden layers, each with fewer nodes. We design the encoder of XVAE with a total of 2 layers prior to the latent embedding. In the first hidden layer, the dimension of each input entity is reduced individually. In the second hidden layer, input entities get fused into a combined layer. The latent embedding (red) represents the bottleneck of the XVAE with the minimum number of nodes. The decoder reversely mirrors the layer structure of the encoder, with the final layer featuring the same number of nodes as the input layer as it attempts to reconstruct (*X*_1_^*′*^, *X*_2_^*′*^) the original input from the latent embedding.

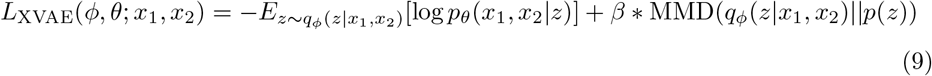

, where *q*_*ϕ*_(*z*|*x*_1_, *x*_2_) encodes the latent space as a probability distribution over the input variables (parameterised by *ϕ*) and *p*_*θ*_(*x*_1_, *x*_2_|*z*) encodes the reconstruction of input variables as a probability distribution over the latent space (parameterised by *θ*). Following the originally proposed implementation, we use maximum mean discrepancy (MMD) as a regularization term to constrain the latent distribution *q*_*ϕ*_ to be a standard Gaussian distribution, balanced by the constant beta (*β*), which is set to 1 for all experiments. A more detailed description on autoencoders, as well as the XVAE architecture and training procedure can be found in the Supplementary Methods.

### 2.4 Multi-omics deconfounding models

Here, we will first describe in section 2.4.1 the use of linear regression for confounder correction and PCA for dimensionality reduction, which we deem the “baseline model” due to their wide popularity. Then, we outline in section 2.4.2 - 2.4.5 the four XVAE-based deconfounding models proposed in this study. Throughout this section we denote the confounder value of a single data point as *c*.

#### 2.4.1 Baseline model: linear regression and PCA (LR+PCA)

Under the assumption that the effects of one or multiple confounders are linearly additive to the true signal of a feature, we build a linear regression (LR) model for the confounders against each mRNA or DNAm feature and then take their residuals as adjusted features. Subsequently, the adjusted features from the two data types are concatenated and their dimensionality is reduced via PCA (LR+PCA). We select the top 50 PCs explaining most of the variance of data to keep the embedding size identical to that of every XVAE-based model. The 50 PCs explaining the most variance of data are considered for the final clustering, for which KMeans with 10 random initialisations is applied.

#### 2.4.2 Conditional X-shaped Variational Autoencoder (cXVAE)

Conditional variational autoencoder (cVAE) [25] is a semi-supervised variation of VAE, which originally aims to fit the distribution of the high-dimensional output as a generative model conditioned on auxiliary variables. Lawry et al. proposed to achieve deconfounding through a cVAE incorporating confounding variable information as auxiliary variables [17]. We extend this initial idea to be able to handle multi-omics data by replacing the originally proposed VAE with the XVAE model, resulting in a conditional X-shaped variational autoencoder (cXVAE) architecture (Figure 3A). We tested the integration of confounders at different levels of the cXAVE, including the input layer, the hidden layer that fuses multiple inputs, and the embedding. More details on cXVAE implementations can be found in the Supplementary Methods.

**Figure 3.**
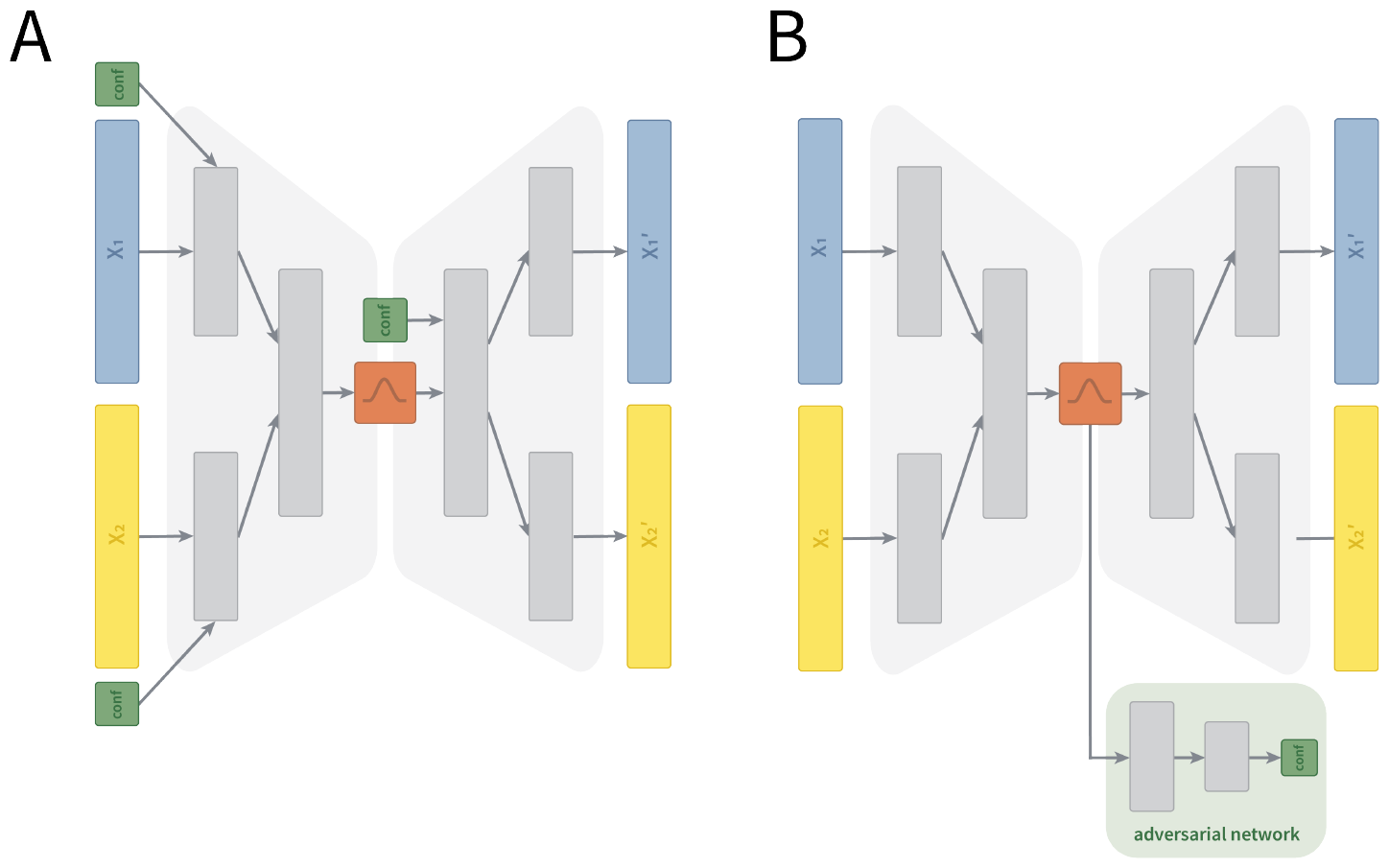
Schematic representation of (A) conditional variational autoencoder (cVAE) and (B) adversarial deconfounding XVAE (adv-XVAE). (A) Depicts the cXVAE implementation termed *input + embed* due to the addition of confounders (green) in the first layer of the encoder and decoder. (B) Depicts the adv-XVAE implementation termed *multiclass* due to the usage of only a single supervised adverarial network (light green) trained to predict confounders (green) using a multiclass prediction loss. *X*_1_ and *X*_2_ are the two omics dimensions, namely gene expression and DNA methylation, while *X*_1_^*′*^ and *X*_2_^*′*^ denote their respective reconstruction. More details and visualisations of other implementation can be found in the Supplementary Material.

#### 2.4.3 X-shaped Variational Autoencoder with adversarial training (adv-XVAE)

The adversarial deconfounding autoencoder proposed by Dincer et al. [7] follows the idea of training two networks simultaneously - an autoencoder to generate a low dimensional embedding and an adversary multi-layer perceptron (MLP) trained to predict the confounder from said embedding (Figure 3B). By adversarially training the two networks, i.e. the autoencoder aims to generate an embedding which can not be used for confounder prediction by the MLP, it aims at generating embeddings that can encode biological information without encoding any confounding signal. As the original framework can only handle a single data type, we adapt it to work with multi-omics input by replacing its autoencoder with XVAE architecture. Details on architecture and training procedure of adv-XVAE can be found in Supplementary Methods.

#### 2.4.4 X-shaped Variational Autoencoder with deconfounding regularization (cr-XVAE)

Augmenting the loss function of deep learning models is an effective way to impose restrictions on the model or enforce learning of specific patterns. As an example, studies focused on disentangling the often highly correlated latent space of autoencoders impose constraints on the correlation between latent features by adding a penalty term to the loss function [18]. Inspired by this idea, we formulate a deconfounding regularization term aiming to reduce the degree of correlation between latent features and confounders. The regularized loss function becomes:

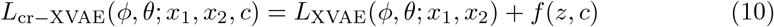

, where *f* (*z, c*) denotes the joint association between latent features and confounders. More specifically, we choose two different association measurements, Pearson correlation and mutual information. Because Pearson correlation ranges from −1 (negatively correlated) to 1 (positively correlated) and both indicate strong relationship, we regularize only the magnitude of correlation by two methods, taking its absolute value or squared value. Because the confounder distribution needed for mutual information is usually unknown, we implement two methods to approximately compute mutual information as loss function, with differentiable histogram or kernel density estimate.

#### 2.4.5 Feature selection by removing correlated latent features (XVAE+FS)

The removal of latent features correlated with confounders comes from the idea of *post hoc* interpretation of latent features [10]. To identify confounded latent features, we calculate the Pearson correlation between each latent variable and the confounder. For determining the threshold indicating which latent features are being removed from further analyses, we test two different approaches:

1. *p-value cutoff* - the p-value of the Pearson correlation indicates the probability that the computed correlation is smaller than a random correlation between uncorrelated datasets. Latent features with a p-value *<* 0.05 are excluded from analyses.

2. *absolute correlation coefficient* - Pearson correlation measures the linear relationship between two variables. Latent features exhibiting an absolute Pearson correlation of more than 0.3 (weak correlation) or 0.5 (strong correlation) are excluded.

### 2.5 Consensus clustering

Different from the baseline linear regression model which adopts KMeans on the deconfounded features for clustering, we apply consensus clustering on the latent features of each VAE-based deconfounding model. Here, consensus clustering takes the advantage of random sampling in a VAE and it constructs a consensus matrix *A* ∈ ℝ^*n×n*^ from the individual clustering of each embedding sampled from the latent distributions [16]. We perform consensus clustering on the embeddings of the entire sample set, generating 50 embedding matrices, on each of which a k-means clustering is conducted. The values in the consensus matrix indicate the fraction of times that two data points are assigned to the same cluster in those 50 clustering solutions. Subsequently, each value is divided by the total number of clusterings (50), resulting in the range [0,1], where 0 means the two corresponding samples are never clustered together while 1 means they are always in the same cluster. Finally, a spectral clustering is performed on the consensus matrix *A* to derive a stable clustering of the patients.

### 2.6 Evaluation metrics

We apply each of the aforementioned models to the artifically confounded multi-omics dataset described in section 2.1 and 2.2. Every model is evaluated in terms of their XVAE reconstruction accuracy, measured as the relative reconstruction error of inputs, their clustering stability, evaluated by the dispersion score of consensus clustering (CC), and deconfounding capabilities for clustering, estimated by calculating the Adjusted Rand index (ARI) for true (cancer types) and confounder labels.

#### 2.6.1 XVAE reconstruction accuracy

Model training is monitored through inspection of the validation loss. To evaluate reconstruction quality of the trained XVAE model, we compute the L2 relative error (RE) between the original input (*x*) and reconstructed data (*x*^*′*^) for (i) each data type individually:

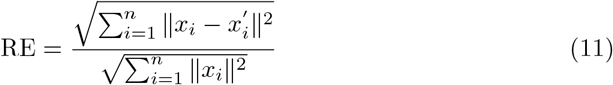

, as well as (ii) for the combined data types:

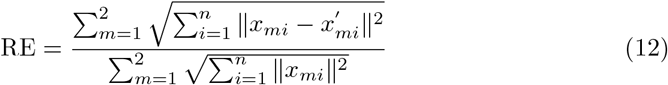

, where *m* = 1, 2 indicates the two data types.

#### 2.6.2 Clustering stability

Before assessing how well each model can derive a meaningful clustering, we want to first check if a model can stably cluster the samples. To achieve this goal, we employ the dispersion score to measure the internal stability of consensus clustering based on its consensus matrix *A*:

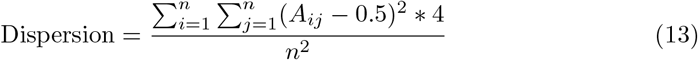

The dispersion score ranges from 0 to 1, where 1 shows a perfect stability that every value in *A* is either 0 or 1, i.e. no confusion among the clusterings, and the lower the less consensus among the clusterings.

#### 2.6.3 Deconfounding capabilities

We compare our clustering with two different labels, the true one, namely cancer types, and the confounder. An ideal model should deconfound the features sufficiently while keeping the meaningful information for obtaining the true clustering. In other words, we expect a model with high ARI with the true label and low ARI with the confounder label. The association between true patient label and clusters obtained when modelling the original (unconfounded) data represents the best achievable clustering, with ARI value converging towards 1.

Similar to ARI, we compute another external clustering metric, the normalized mutual information (NMI), which measures the dependence between two clusterings. As it only shows complementary information to ARI, we record the NMI of every clustering model in Supplementary Table 1.

### 2.7 Implementation

For better stability and generalization, we train each model 50 times using i) randomly sampled training and validation sets with a ratio of 80:20 and ii) different seed of randomization.

### 2.8 Software

All of the models described in this study are built in Pytorch Lightning [9] and trained using the GPU units RTX 2080 Ti 11GB.

## 3 Results

### cXVAE outperforms other considered deconfounding strategies in the presence of a single confounder

We simulated different types of confounding effects - linear, non-linear (squared), and categorical - on the multi-omics TCGA pan-cancer dataset to benchmark a total of four deconfounding frameworks, namely XVAE with Pearson correlation feature selection (XVAE+FS), conditional XVAE (cXVAE), adversarial training with XVAE (adv-XVAE), and confounder-regularised XVAE (cr-XAVE) (see Methods for more details). We additionally included two baseline models to compare with: 1) confounder correction with linear regression (LR+PCA) and 2) vanilla XVAE without any deconfounding (XVAE). To estimate the robustness of each method, each model was trained on 50 iterations of randomly sampled training and validation data (80:20 split) and random seed initialization.

All proposed deconfounding approaches were able to correct for a linear confounder, as denoted by the high ARI for true clustering and low ARI for confounder clustering (Table 1). Performances started to decline for non-linear confounding problems, with cXVAE clearly outperforming other strategies. For non-linear confounders we noted large ARI for confounder clustering across all strategies and simulation setups. This illustrates that, while good clustering performance for true labels were achieved, the full removal of unwanted signal was not easily achievable for all the models. Categorical confounding was perceived to be the most difficult, with all models except cXVAE exhibiting a high decrease in true clustering performance. Notably, cr-XVAE and XVAE+FS were able to remove artificial confounders completely, however at the cost of simultaneously removing true clustering signal. adv-XVAE, which in theory should be a strategy well suited to deal with categorical problems, fails to consistently remove the categorical confounding effect. In general we noted a decline of reconstruction accuracy of models with increasing complexity of the confounder simulations.

**Table 1:** Overview performances of deconfounding strategy for single confounder simulations. Values are displayed as mean *±* standard deviation of 50 runs with different parameter initialisation and randomly sampled training and validation data. Models on the first column indicate the following deconfounding strategies and implementations thereof: linear regression followed by principal component analysis and KMeans clustering (LR+PCA), vanilla XVAE without any deconfounding (XVAE), XVAE with feature selection in the form of removing correlated latent features (XVAE+FS, correlation cutoff = 0.5), conditional XVAE (cXVAE, input + embedding), adversarial training with XVAE (adv-XVAE, multiclass MLP), confounder-regularised XVAE (cr-XAVE, squared correlation regularisation). Reconstruction error: relative error in the reconstruction of X1 and X2 weighted eqally; CC dispersion: consensus clustering agreement over 50 iterations; True clustering: adjusted rand index (ARI) of consensus clustering derived clusters with True label labels; Confounder clustering: ARI of consensus clustering derived clusters with simulated confounder labels.

| linear |  |  |  |  |
| --- | --- | --- | --- | --- |
|  | Reconstruction error | CC dispersion | Clustering performance (ARI) |  |
|  |  |  | True label (cancer type) | Confounder |
| LR+PCA | - | - | 0.692 | 0.001 |
| XVAE | 0.246 (± 0.004) | 0.844 (± 0.045) | 0.506 (± 0.116) | 0.151 (± 0.057) |
| XVAE+FS | 0.245 (± 0.004) | 0.860 (± 0.028) | 0.571 (± 0.092) | 0.008 (± 0.007) |
| cXVAE | **0.234 (± 0.003)** | **0.935 (± 0.023)** | **0.712 (± 0.055)** | **0.001 (± 0.001)** |
| adv-XVAE | 0.245 (± 0.004) | 0.901 (± 0.032) | 0.568 (± 0.070) | 0.093 (± 0.051) |
| cr-XVAE | 0.244 (± 0.003) | 0.873 (± 0.028) | 0.598 (± 0.074) | 0.004 (± 0.004) |
| non-linear |  |  |  |  |
|  | Reconstruction error | CC dispersion | Clustering performance (ARI) |  |
|  |  |  | True label (cancer type) | Confounder |
| LR+PCA | - | - | 0.391 | 0.215 |
| XVAE | 0.236 (± 0.003) | 0.826 (± 0.042) | 0.307 (± 0.141) | 0.297 (± 0.071) |
| XVAE+FS | 0.236 (± 0.003) | 0.805 (± 0.040) | 0.411 (± 0.142) | 0.138 (± 0.078) |
| cXVAE | **0.227 (± 0.002)** | **0.908 (± 0.031)** | **0.646 (± 0.079)** | **0.076 (± 0.074)** |
| adv-XVAE | 0.238 (± 0.004) | 0.942 (± 0.025) | 0.568 (± 0.049) | 0.194 (± 0.006) |
| cr-XVAE | 0.235 (± 0.002) | 0.852 (± 0.043) | 0.478 (± 0.129) | 0.154 (± 0.042) |
| categorical |  |  |  |  |
|  | Reconstruction error | CC dispersion | Clustering performance (ARI) |  |
|  |  |  | True label (cancer type) | Confounder |
| LR+PCA | - | - | 0.150 | 0.071 |
| XVAE | 0.216 (± 0.003) | 0.762 (± 0.055) | 0.330 (± 0.125) | 0.048 (± 0.088) |
| XVAE+FS | 0.216 (± 0.003) | 0.787 (± 0.040) | 0.361 (± 0.100) | 0.010 (± 0.023) |
| cXVAE | **0.210 (± 0.002)** | **0.911 (± 0.033)** | **0.664 (± 0.070)** | **0.001 (± 0.001)** |
| adv-XVAE | 0.217 (± 0.002) | 0.764 (± 0.058) | 0.240 (± 0.188) | 0.156 (± 0.084) |
| cr-XVAE | 0.216 (± 0.003) | 0.813 (± 0.034) | 0.368 (± 0.101) | 0.001 (± 0.001) |

Figure 4 visualizes the deconfounding behaviour of cXVAE for categorical confounding. With an exception to THCA (thyroid carcinoma), all classes were strongly confounded prior to cXVAE application (Figure 4, left). After model training, confounding classes are homogeneously mixed and clustering occurs with respect to true cancer types (Figure 4, right). In summary, across all confounder simulations, cXVAE clearly outperformed other deconfounding strategies in terms of clustering accuracy, deconfounding power, and model robustness. The ARI on true clustering obtained by cXVAE in all three scenarios reached around 0.7, which is very close to the performance of the vanilla XVAE on unconfounded data (0.731, see details in Supplementary Table 2).

**Figure 4.**
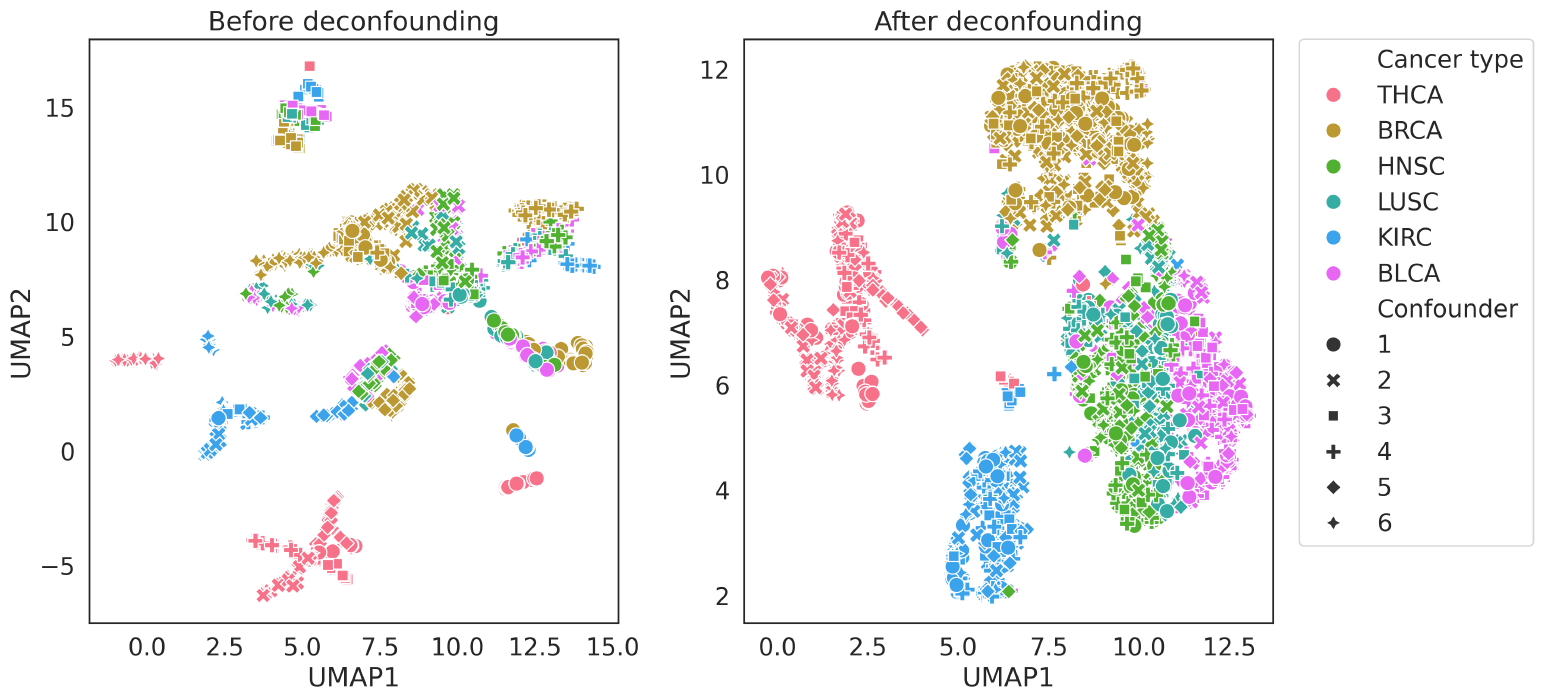
Deconfounding behaviour of cXVAE. Dimensionality reduction (UMAP) plot of categorically confounded data before and after application of cXVAE. Marker colors indicate the true label labels (i.e. TCGA cancer types), while marker shapes indicate the six classes (1-6) of the confounder (see section 2.2).

A more detailed summary of the performances of each model can be found in Supplementary Table 1.

While Table 1 depicts the best performing implementation of each deconfounding model, we tested a number of possible implementations (see Methods), which we observed to have a notable impact on model performance (Supplementary Table 2). Therefore, we provide design recommendations for each deconfounding strategy in the Supplementary Results.

### cXVAE is easily extendable to handle multiple confounders of mixed types

In a realistic setting datasets can be confounded by multiple confounders with different biasing effects. In an attempt to investigate how well deconfounding strategies can handle more than one confounder, we simulated the parallel presence of three confounders of different effect, namely linear, non-linear, and categorical (Table 2).

**Table 2:**
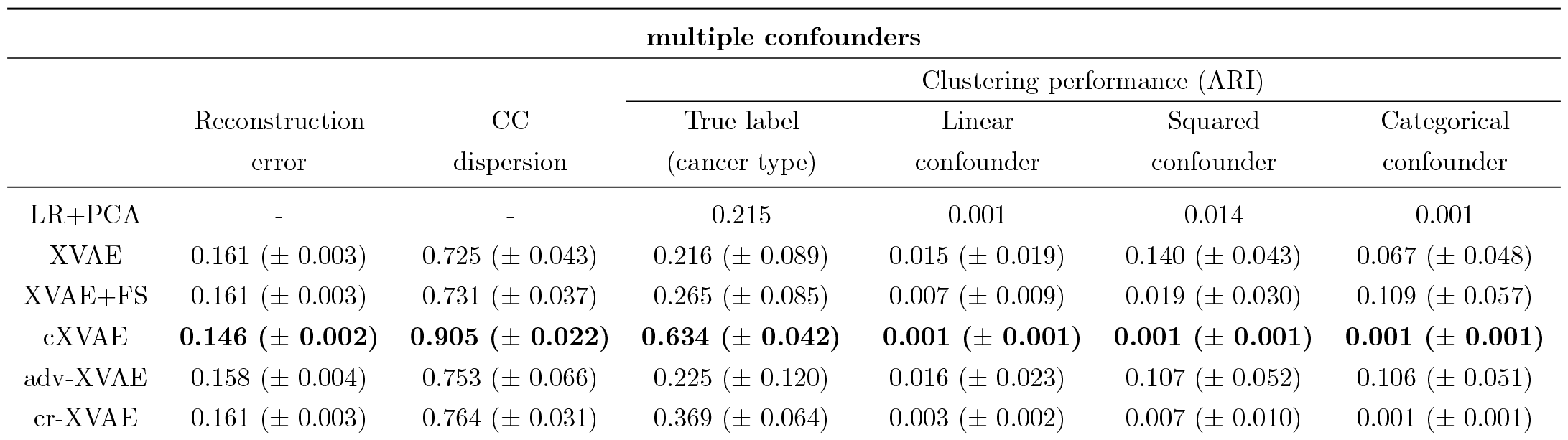
Overview performances of deconfounding strategy in the presence of multiple confounders. Values are displayed as mean *±* standard deviation of 50 runs with different parameter initialisation and randomly sampled training and validation data. For a detailed description of columns and models, please refer to Table 1.

In line with our observations with the single confounder simulations, cXVAE outperformed other models in terms of true clustering accuracy and de-confounding efficiency. While also other strategies like XVAE+FS, cr-XVAE, or LR+PCA were able to successfully remove all three simulated effects, they achieved this at the cost of true signal. adv-XVAE failed to fully remove confounders, while also showing very low true clustering accuracy and can therefore be considered unsuitable for the task. We also noted that the decline in reconstruction accuracy with increasingly complex confounding situations is even more pronounced in multiple confounder settings.

### cXVAE is able to retrieve biology-driven clustering from confounded data

To illustrate the deconfounding capabilities of cXVAE, the model that outperformed others across all four evaluation metrics in various confounding scenarios, we examined the clustering results obtained on the TCGA dataset involving categorical confounders (Figure 4, right). The UMAP plot of latent features clearly showed that BRCA (breast invasive carcinoma), THCA (thyroid carcinoma), and KIRC (kidney renal clear cell carcinoma) were well clustered by cXVAE, while BLCA (bladder urothelial carcinoma), LUSC (lung squamous cell carcinoma), and HNSC (head and neck squamous cell carcinoma) were still entangled. In summary, we found this behaviour to be in line with the pathological and physiological differences between these cancer types. BLCA arises from urothelial cells in the transitional epithelium, which can change from cuboidal to squamous form when stretched. Furthermore, squamous differentiation is by far the most common histological variant of urothelial carcinoma [14], indicating a close relationship between urothelial carcinoma and squamous cell carcinoma. Apart from BLCA, the overlap in clustering of LUSC and HNSC can be directly explained by their common origin of squamous cells, while BRCA, THCA, and KIRC are all carcinoma related to glandular cells [13]. Supporting the validity of our obtained cXVAE clustering, other multi-omics pan-cancer studies utilising stacked variational autoencoders [27], penalized matrix factorization [12], or supervised VAE [31] have retrieved similar cancer type clustering.

## 4 Discussion

In this study, we addressed the possible harm of ignoring or inadequately handling confounders to clustering samples with (multi-)omics measurements. In epidemiology, a confounder is a variable that can effect the result of a study because it is related to both the exposure and the outcome being studied. Here, we extended the definition to unsupervised models for disease subtyping to indicate variables that can distort the relationship between inferred or predicted cluster membership and disease.

Extensive simulation revealed that cXVAE stands out as a versatile and accurate deconfounding approach. The applicability of conditional autoencoder to biological data to e.g. correct for batch effects [17] or disentangle confounders in fMRI [28] or microRNA data [30] has been shown before. However, by merging the principles of a conditional autoencoder with the framework of an autoencoder specifically tailored for the integration of multi-omics data, our research charts new frontiers in the domain of deconfounded patient stratification.

While adversarial training may offer an alternative flexible deconfounding approach, we confirm that optimization of model hyper-parameters is challenging [2]. Instability may become more pronounced in the presence of multiple confounders. This can be explained by the fact that adversarial networks were trained separately for each confounder, sequentially adding extra terms to its objective function (see Supplementary Table 2).

In the literature, a statistical correlation loss has been proposed to replace the adversarial prediction loss in a adversarial training model [1], resembling our cr-XVAE model. The difference is that cr-XVAE directly computes the correlation between the VAE embedding and the confounder without an additional adversarial network. We implemented Pearson correlation and mutual information as the regularization term of cr-XVAE but other association measures could also be adopted, e.g. Spearman correlation and cosine similarity. In the case of multiple confounders, it is also possible to weigh their associations differently in the loss function to balance deconfounding strength.

The identification of disease subtypes requires performing a clustering algorithm at some point. Even though iterative training of the clustering in a joint autoencoder loss function can overcome inconsistencies between training and downstream clustering performance [8, 19, 21], we chose for a decoupled strategy. This was to 1) avoid having too many terms in loss function to confuse training, and 2) reduce computation time and initialization settings with iteratively training clustering in a joint loss function. Consensus clustering furthermore has several advantages in data science including robustness, stability, interpretability and flexibility, as it can be applied to various types of data and clustering algorithms.

It remains a daunting task to generate data that adequately reflect the complexity of real-life cases. Therefore, one needs to be aware that simulations of confounders always represent simplifications of real observable effects. While this study is limited to the use of two data types, in principle the XVAE design utilised allows the integration of heterogeneous data from many more sources simultaneously. Additionally, since all evaluated deconfounding strategies share the same XVAE design as a foundation, we anticipate consistent training time and performance across models when scaling up dimensions.

## 5 Conclusion

In this study, we presented four VAE-based multi-omics clustering models and their variations, following different deconfounding strategies. Their clustering and deconfounding performance was evaluated and compared with baseline models on the multi-omics pancancer dataset from TCGA with artificailly generated confounding effects. The results showed both the necessity to adjust for confounders and that our novel models, cXVAE in particular, can effectively deal with the confounding effects and obtain the biologically meaningful clustering. We demonstrate that our multi-omics deconfounding VAE clustering models have big potential in delivering accurate patient subgrouping or disesase subtyping, ultimately enabling better personalised healthcare.

## Data and code availability

Our code is available at https://github.com/ZuqiLi/Multi-view-Deconfounding-VAE. The simulated data generated in the course of this study are available at Zenodo (https://doi.org/10.5281/zenodo.10458941).

## Acknowledgements

The authors thank the supporters of this study, namely the European Union’s Horizon 2020 research and innovation program under the Marie Sklodowska-Curie grant agreement No. 860895 TranSYS. Furthermore we thank the members of the Computational Population Biology group at Erasmus Medical Center for their critical and creative input to this work.

## Author contributions statement

Z.L. and S.K. designed, planned, carried out the practical aspects of this study, and wrote the manuscript. G.V.R. offered mentoring and scientific input throughout the work process. G.V.R., E.S., K.v.S., and V.MdS. provided critical revision of the manuscrip All authors approve of the final manuscript.

