## Supplementary Material for "Novel multi-omics deconfounding variational autoencoders can obtain meaningful disease subtyping"

†Authors contributed equally *Corresponding author

### Supplementary Results

#### Differences in model implementations have significant impact on deconfounding performance

Each of the deconfounding strategies compared in this study have a number of possible implementations for both single and multiple confounder simulations (Supplementary Table 2), which we observed to have a major impact on model performances. Hence, in this section, we sequentially discuss each strategy, highlighting the most notable differences between implementations and providing readers with recommendations on designing similar frameworks.

**XVAE with feature selection (XVAE+FS)** - the central question when conducting feature selection by removing latent features correlated with unwanted confounders is the correlation cutoff employed. We compared the use of two different correlation cutoffs (Pearson correlation $> 0.5$, correlation $> 0.3$) and using a significant p-value as cutoff (p-value $<0.05$). While using absolute correlations as threshold removed up to 30% of latent features, using the p-value commonly removed >90% of features, rendering the latent space too sparse to use.

**Conditional XVAE (cXVAE)** - cXVAE may vary regarding which layer of the autoencoder confounders are added, and whether they are added in both the encoder and decoder of the autoencoder. We found that in general it does not affect the deconfounding performance of the network in which layer of the encoder confounders are added (Supplementary Table 2, *input + embedding*, *fused + embedding*). We hypothesised that it may suffice to only add confounders to the encoder, however we found that a decoder lacking confounders (*input*) performs significantly worse. Conversely, adding the conditional variable to only the decoder results in good deconfounding performances (*embed*).

**Adversarial training (adv-XVAE)** - while the possibilities to design adversarial training strategies are countless we found it challenging to hyperparameter tune and balance the regularisation terms to simulatenously achieve good reconstruction and satisfactory deconfounding. Bahrami et al. [1] proposed a variation in which a simplified loss is backpropagated solely through the encoder of the network, to reinforce training (*scGAN*). However, also this modified version could not deliver consistent results. We attempted to extend the original model to accommodate multiple confounders by incorporating one adversarial network for each confounder present (*multinet MLP*). Contrary to expectations however, this implementations failed to improve adversarial training performance for the multiple confounder simulations (Supplementary Table 3).

**Confounder regularised XVAE (cr-XVAE)** - the regularisation term added for our deconfounding regularisation strategy largely determines the deconfounding properties. As a general guideline, Pearson correlation accounts for linear associations whereas mutual information can capture higher-level dependence. Because we are only interested in the strength of correlation, not the sign, the correlation coefficient was converted into absolute value or squared, leading to slightly different performances (Supplementary Table 2). Meanwhile, we conducted two different implementations for mutual information as loss function to approximate latent feature distribution, one based on differentiable histogram and the other on kernel density estimate. For both methods, the regularisation quality relies on how good the approximation is.

#### Deconfounding approaches should be motivated by the nature of confounders

Our results revealed that methods differentiate themselves regarding the trade-off they achieve between removing confounding signal and maintaining true patient clustering. According to this, we categorised all surveyed deconfounding approaches in either *aggressive* or *preserving* methods.

*Aggressive methods*, e.g. removing latent features correlated with confounders, or enforcing deconfounding in the objective function, tend to efficiently remove confounders, however at the cost of biological signal. While this stringent approach resulted in intermediate true clustering accuracies in our study, the remaining true signal will likely contain strong true positives findings.

*Preserving methods*, including cXVAE and adversarial training, tend to not remove confounders at all costs, which generally yielded higher true clustering accuracies, but also a higher fraction of deconfounding signal remained in the data.

We argue that aggressive methods are best applied in the case of technical confounders, like batch effects, in which not removing all of the confounding signal may easily lead to spurious biological conclusions [2, 3]. Preserving methods, on the other hand, are interesting for removing biological confounders, e.g. sex or age effects, as applying a too stringent method may lead to removing too much (weak) biological signal of interest. However, this comes at the risk of findings containing more false positives, so thorough validation of results is advised.

Notably, we observed that simply training an autoencoder on confounded data may already lead to deconfounding, which is grounded in the inability of the autoencoder to fully model the confounder. Nevertheless, the obscuring effect of confounders on the true clustering can not be removed by a vanilla autoencoder, underlying the need for extraneous deconfounding approaches.

### Supplementary Methods

#### Visualization of confounder simulations

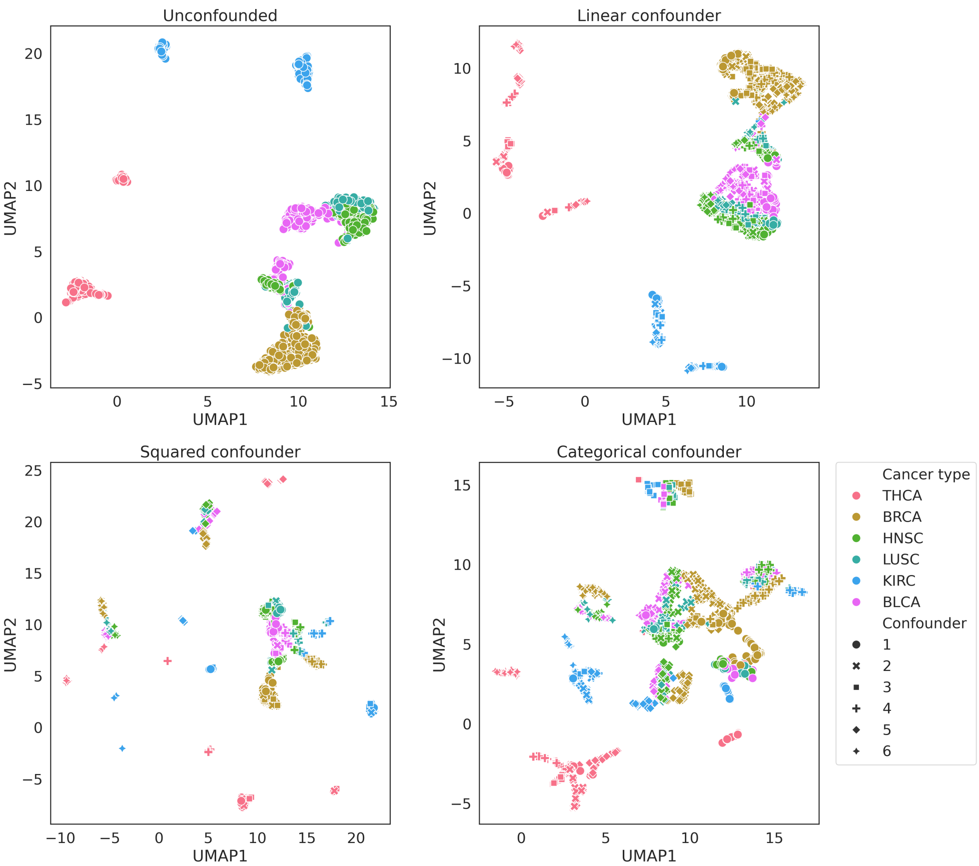

Supplementary Figure 1: Dimensionality reduction (UMAP) plots of unconfounded (upper left), linear confounded (upper right), non-linear confounded (lower left), and categorical confounded (lower right) data. Colors indicate the true labels (i.e. TCGA cancer types), while shapes indicate the six different simulated confounders

#### Variational autoencoder for data integration (XVAE)

Autoencoders (AE) are unsupervised neural networks consisting of an encoder and a decoder part. While the encoder ($E_\phi$, parameterised by $\phi$) maps the high dimensional input datapoint ($x$) into a lower dimensional latent embedding ($z$), the decoder ($D_\theta$, parameterised by $\theta$) attempts to reconstruct the original input from said embedding ($x' = D_\theta(E_\phi(x))$).

The network is then trained by trying to minimize the error between original input and reconstruction, quanitfied by the mean squared error:

$$L_{\mathrm{AE}}(\phi, \theta; x) = \|x - x'\|^2 = \|x - D_\theta(z)\|^2 = \|x - D_\theta(E_\phi(x))\|^2 \tag{1}$$

Standard autoencoders have been demonstrated shortcomings in terms of generative abilities and the tendency to overfit input data due to their lack of regularization. These limitations are addressed by a popular variant of autoencoders, termed variational autoencoders (VAE), which encodes input variables as a probability distribution ($q_\phi(z|x)$) over the latent space rather than as a fixed value [4]. For the sake of easy computation and interpretation, the typical choice is to assume the latent distribution $q_\phi(z|x)$ as a standard Gaussian distribution The unsupervised encoder-decoder structure allows (variational) autoencoders to act as dimensionality reduction and clustering tools, capable of accommodating input data from different sources in a joint low-dimensional embedding, ultimately making them a popular tool for data integration. Following the originally proposed implementation, we used a combined loss function of the Mean Squared Error (MSE) for scoring the reconstruction loss and Maximum Mean Discrepancy (MMD) as regularization term, balanced by the constant beta ($\beta$), which was set to 1 for all experiments:

$$L_{\mathrm{VAE}}(\phi, \theta; x) = -E_{z \sim q_\phi(z|x)}[\log p_\theta(x|z)] + \beta * \mathrm{MMD}(q_\phi(z|x)||p(z)) \tag{2}$$

For the activation function, we choose the Leaky Rectified Linear Activation Function (LeakyReLU) [5] in all layers except the output layer, which employs a sigmoid activation function. We are able to use the sigmoid function due to the normalisation of every mRNA and DNAm feature to the range $[0, 1]$ during preprocessing. To enhance training stability, batch normalisation layers are added to the encoder. We train every model for a fixed period of 150 epochs using an Adam optimiser [6] and a batch size of 64. To derive an optimal architecture for the XVAE base model, we carry out a hyperparameter search assessing the combination of (1) the number of nodes in hidden layers and latent embedding, (2) the rate of dropout layers, and (3) the weight initialisation.

#### cXVAE: Conditional XVAE

While traditionally auxiliary variables are added in the encoder and decoder, in theory this can be varied. To investigate the impact on deconfounding attributable to when confounders are added to the model, we created several variations of the cXVAE architecture (Supplementary Figure 2):

1. *input* - in the encoder confounders are concatenated with the input vector; no addition of confounders in the decoder
2. *embed* - in the decoder confounders are concatenated with the latent embedding; no addition of confounders in the encoder
3. *input + embedding* - in the encoder confounders are concatenated with the input vector; in the decoder confounders are concatenated with the latent embedding
4. *fused + embedding* - in the encoder confounders are concatenated in the second hidden layer, where the fusion of data types occurs; in the decoder confounders are concatenated with the latent embedding

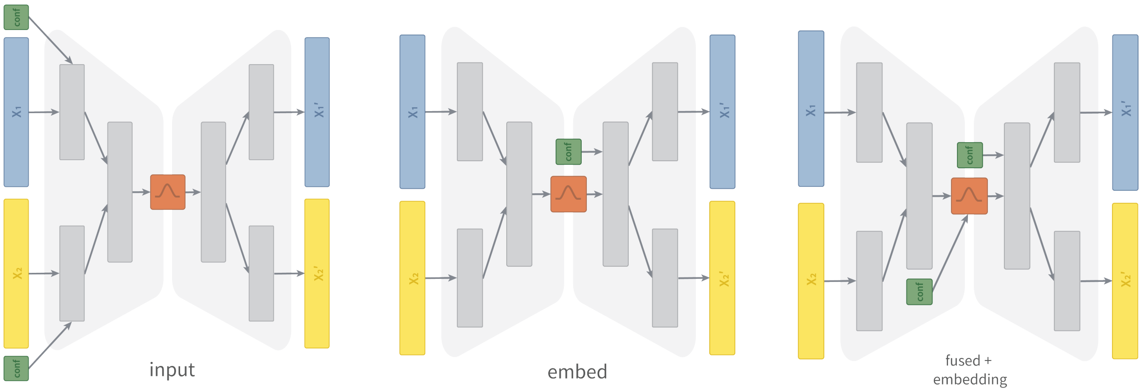

Supplementary Figure 2: Overview on the different cXVAE assessed in this study.

#### adv-XVAE: XVAE with adversarial training

As an adversarial network we implement a simple MLP with two layers. The first layer takes the generated embedding of the XVAE model as input and applies ReLU activation to condense it to 10 features. The output of the second layer features the number of confounders to be predicted and uses the activation function fitting for predicting respective confounder; i.e. uses ReLU in case of continuous confounders, a sigmoid activation function in case of binary confounders, and softmax activation for multiple categorical confounders.

The training procedure can be broken down in three steps:

(i) Pre-train the XVAE: we pre-trained the XVAE model for 5 epochs optimising equation 9.

(ii) Pre-train the adversarial network: we used the latent embedding initialized in (i) to pre-trained the adversarial network for 5 epochs using a loss function fitting the numerical type of confounder; mean squared error for continuous, binary cross entropy for binary, and crossEntropy for categorical confounders, i.e. $L_{\mathrm{adv}}(h_\nu(x), c)$ where $h_\nu$ denotes the adversarial network parameterized by $\nu$.

(iii) Joint adversarial training: joint training is unique in its attempt to minimize the XVAE loss (equation 9) while simultaneously maximizing the adversarial network loss through a combined loss function:

$$L_{\mathrm{adv-XVAE}}(\phi, \theta, \nu; x, c) = L_{\mathrm{XVAE}}(\phi, \theta; x) - \lambda L_{\mathrm{adv}}(h_v(x), c)] \qquad (3)$$

Training is carried out in a ping-pong fashion, where firstly all weights of the adversarial network are frozen and the autoencoder trained for one epoch optimizing equation 2, followed by the freezing of autoencoder weights and training of the adversary network for an entire epoch to minimize the adversarial network prediction loss $L_{\mathrm{adv}}$.

Depending on how the adversarial architecture is designed in the case of multiple confounders present, we differentiated two major variations the above described base model: multiclass MLP and multinet MLP.

In *multiclass MLP* adversarial networks confounders are concatenated to a combined label, effectively creating a (mixed) multiclass classification problem. To successfully predict labels of different numeric nature, instead of a single final layer, multiclass MLP feature up to three final layers, each with different activation functions (ReLU for numerical confounders, Sigmoid for binary categorical confounders, softmax for categorical confounders with more than one class).

*Multinet MLP*, on the other hand, possesses an individual adversarial MLP for each confounder. The adversarial loss $L_{\text{adv}}$ becomes the sum of losses of these individual networks.

Instead of augmenting the architecture of their *scGAN* model, Bahrami et al. [1] proposed changes to the training procedure. While following the same initial strategy of separately pre-training the autoencoder and adversarial MLP, they simplify the overall objective function (equation 3) by omitting to autoencoder loss and instead only train the encoder by maximizing the adversarial prediction loss $L_{\text{adv}}$.

### Supplementary Tables

**linear confounder**

| | | reconstruction error | | | CC | Internal clustering | | True clustering | | Confounder clustering | |
|---|---|---|---|---|---|---|---|---|---|---|---|
| | | X1 | X2 | X1,X2 | dispersion | Silhouette | DB index | ARI | NMI | ARI | NMI |
| XVAE | mean | 0.303 | 0.222 | 0.246 | 0.844 | 0.120 | 2.114 | 0.506 | 0.594 | 1.51E-01 | 2.02E-01 |
| | std | 0.010 | 0.003 | 0.004 | 0.045 | 0.013 | 0.118 | 0.116 | 0.093 | 5.71E-02 | 7.13E-02 |
| XVAE + FS | mean | 0.302 | 0.222 | 0.245 | 0.860 | 0.131 | 2.076 | 0.571 | 0.686 | 7.70E-03 | 1.14E-02 |
| | std | 0.010 | 0.003 | 0.004 | 0.028 | 0.011 | 0.146 | 0.092 | 0.053 | 6.79E-03 | 7.44E-03 |
| cXVAE | mean | 0.286 | 0.212 | 0.234 | 0.935 | 0.133 | 2.243 | 0.712 | 0.786 | 4.82E-04 | 3.11E-03 |
| | std | 0.008 | 0.001 | 0.003 | 0.023 | 0.011 | 0.092 | 0.055 | 0.037 | 5.37E-04 | 7.02E-04 |
| adv-XVAE | mean | 0.302 | 0.221 | 0.245 | 0.901 | 0.143 | 2.013 | 0.568 | 0.666 | 9.34E-02 | 1.28E-01 |
| | std | 0.010 | 0.003 | 0.004 | 0.032 | 0.011 | 0.099 | 0.070 | 0.053 | 5.14E-02 | 6.60E-02 |
| cr-XVAE | mean | 0.300 | 0.221 | 0.244 | 0.873 | 0.129 | 2.133 | 0.598 | 0.715 | 4.18E-03 | 6.97E-03 |
| | std | 0.007 | 0.002 | 0.003 | 0.028 | 0.010 | 0.123 | 0.074 | 0.046 | 3.91E-03 | 3.97E-03 |

**non-linear confounder**

| | | reconstruction error | | | CC | Internal clustering | | True clustering | | Confounder clustering | |
|---|---|---|---|---|---|---|---|---|---|---|---|
| | | X1 | X2 | X1,X2 | dispersion | Silhouette | DB index | ARI | NMI | ARI | NMI |
| XVAE | mean | 0.235 | 0.236 | 0.236 | 0.826 | 0.119 | 2.078 | 0.307 | 0.424 | 2.97E-01 | 3.71E-01 |
| | std | 0.005 | 0.002 | 0.003 | 0.042 | 0.011 | 0.099 | 0.141 | 0.120 | 7.07E-02 | 8.17E-02 |
| XVAE + FS | mean | 0.235 | 0.236 | 0.236 | 0.805 | 0.114 | 2.120 | 0.411 | 0.526 | 1.38E-01 | 1.84E-01 |
| | std | 0.005 | 0.002 | 0.003 | 0.040 | 0.011 | 0.111 | 0.142 | 0.123 | 7.80E-02 | 9.75E-02 |
| cXVAE | mean | 0.223 | 0.229 | 0.227 | 0.908 | 0.124 | 2.185 | 0.646 | 0.713 | 7.59E-02 | 1.06E-01 |
| | std | 0.005 | 0.001 | 0.002 | 0.031 | 0.012 | 0.070 | 0.079 | 0.076 | 7.36E-02 | 9.87E-02 |
| adv-XVAE | mean | 0.240 | 0.237 | 0.238 | 0.942 | 0.147 | 1.918 | 0.568 | 0.635 | 0.194 | 0.249 |
| | std | 0.010 | 0.002 | 0.004 | 0.025 | 0.012 | 0.090 | 0.049 | 0.032 | 0.006 | 0.007 |
| cr-XVAE | mean | 0.235 | 0.235 | 0.235 | 0.852 | 0.121 | 2.092 | 0.478 | 0.575 | 1.54E-01 | 2.08E-01 |
| | std | 0.005 | 0.001 | 0.002 | 0.043 | 0.011 | 0.094 | 0.129 | 0.092 | 4.15E-02 | 5.24E-02 |

Supplementary Table 1: **Overview performances of deconfounding strategy for single confounder simulations**. Values are displayed as mean (mean) and standard deviation (std) of 50 runs with different initialisation. Models on the first column indicate the following deconfounding strategies and implementations thereof: vanilla XVAE without any deconfounding (XVAE), XVAE with feature selection in the form of removing correlated latent features (XVAE+FS, correlation cutoff = 0.5), conditional XVAE (cXVAE, input + embedding), adversarial training with XVAE (adv-XVAE, multiclass MLP), confounder-regularised XVAE (cr-XAVE, squared correlation regularisation). Reconstruction error: relative error in the reconstruction of X1 and/or X2; CC dispersion: consensus clustering agreement over 50 iterations; Internal clustering: ....; True clustering: adjusted rand index (ARI) and Normalized Mutual Information (NMI) of consensus clustering derived clusters with ground-truth labels; Confounder clustering: ARI and NMI of consensus clustering derived clusters with simulated confounder labels.

**categorical confounder**

| | | reconstruction error | | | CC | Internal clustering | | True clustering | | Confounder clustering | |
|---|---|---|---|---|---|---|---|---|---|---|---|
| | | X1 | X2 | X1,X2 | dispersion | Silhouette | DB index | ARI | NMI | ARI | NMI |
| XVAE | mean | 0.205 | 0.224 | 0.216 | 0.762 | 0.085 | 2.379 | 0.330 | 0.431 | 4.85E-02 | 9.12E-02 |
| | std | 0.006 | 0.002 | 0.003 | 0.055 | 0.008 | 0.073 | 0.125 | 0.148 | 8.80E-02 | 1.44E-01 |
| XVAE + FS | mean | 0.205 | 0.224 | 0.216 | 0.787 | 0.084 | 2.327 | 0.361 | 0.473 | 9.53E-03 | 2.02E-02 |
| | std | 0.006 | 0.002 | 0.003 | 0.040 | 0.008 | 0.087 | 0.100 | 0.099 | 2.31E-02 | 4.50E-02 |
| cXVAE | mean | 0.197 | 0.218 | 0.210 | 0.911 | 0.117 | 2.311 | 0.664 | 0.738 | 4.67E-04 | 3.17E-03 |
| | std | 0.003 | 0.001 | 0.002 | 0.033 | 0.011 | 0.084 | 0.070 | 0.060 | 9.62E-04 | 1.34E-03 |
| adv-XVAE | mean | 0.205 | 0.224 | 0.217 | 0.764 | 0.121 | 2.168 | 0.240 | 0.273 | 1.56E-01 | 3.12E-01 |
| | std | 0.003 | 0.001 | 0.002 | 0.058 | 0.008 | 0.050 | 0.188 | 0.203 | 8.40E-02 | 1.60E-01 |
| cr-XVAE | mean | 0.204 | 0.224 | 0.216 | 0.813 | 0.109 | 2.077 | 0.368 | 0.490 | -9.12E-05 | 2.55E-03 |
| | std | 0.005 | 0.002 | 0.003 | 0.034 | 0.011 | 0.091 | 0.101 | 0.098 | 6.09E-04 | 7.96E-04 |

Supplementary Table 2: **Overview performances of deconfounding strategy for single confounder simulations** - continued

Supplementary Table 2: **Performance overview of different implementations of each deconfounding strategy for single confounders**

**linear confounder**

| | | | | | | Clustering performance | | | | | |
|---|---|---|---|---|---|---|---|---|---|---|---|
| | Implementation | Reconstruction error | | | CC | Internal | | True label | | Confounder | |
| | | X1 | X2 | X1,X2 | dispersion | Silhouette | DB index | ARI | NMI | ARI | NMI |
| baseline | XVAE (unconfounded) | 0.438 | 0.278 | 0.31 | 0.957 | 0.128 | 2.17 | 0.731 | 0.821 | 1.14E-04 | 2.60E-03 |
| | KMeans | - | - | - | - | 0.132 | 2.14 | 0.212 | 0.381 | 3.53E-01 | 5.15E-01 |
| | PCA(50) + KMeans | - | - | - | - | 0.189 | 1.66 | 0.211 | 0.381 | 3.54E-01 | 5.17E-01 |
| | LR + PCA + KMeans | - | - | - | - | 0.251 | 1.80 | 0.692 | 0.763 | 1.50E-04 | 2.66E-03 |
| | XVAE | 0.294 | 0.22 | 0.241 | 0.807 | 0.11 | 2.25 | 0.411 | 0.522 | 2.08E-01 | 2.93E-01 |
| XVAE + FS | XVAE (corr 0.3) | 0.294 | 0.220 | 0.241 | 0.843 | 0.133 | 2.12 | 0.561 | 0.634 | 6.54E-03 | 1.13E-02 |
| | XVAE (corr 0.5) | 0.294 | 0.220 | 0.241 | 0.868 | 0.124 | 2.22 | 0.617 | 0.710 | 4.73E-03 | 8.51E-03 |
| | XVAE (pVal 0.05) | 0.294 | 0.220 | 0.241 | 0.653 | 0.160 | 1.66 | 0.261 | 0.354 | 3.38E-02 | 4.96E-02 |
| cXVAE | input | 0.29 | 0.217 | 0.238 | 0.907 | 0.111 | 2.06 | 0.429 | 0.582 | 1.84E-01 | 2.23E-01 |
| | inputEmbed | 0.281 | 0.212 | 0.233 | 0.895 | 0.147 | 2.21 | 0.715 | 0.768 | 4.72E-04 | 3.13E-03 |
| | embed | 0.285 | 0.215 | 0.235 | 0.931 | 0.146 | 2.20 | 0.702 | 0.769 | 4.87E-04 | 3.06E-03 |
| | fusedEmbed | 0.282 | 0.212 | 0.233 | 0.933 | 0.127 | 2.26 | 0.718 | 0.764 | 5.41E-04 | 3.15E-03 |
| adv-XVAE | multiclass | 0.289 | 0.217 | 0.238 | 0.920 | 0.142 | 2.16 | 0.632 | 0.723 | 3.46E-03 | 6.98E-03 |
| | multiclass (1batch) | 0.358 | 0.272 | 0.297 | 0.867 | 0.150 | 1.79 | 0.368 | 0.509 | 1.62E-01 | 3.05E-01 |
| | multiclass (scGAN) | 0.287 | 0.214 | 0.235 | 0.940 | 0.125 | 2.26 | 0.739 | 0.821 | 2.23E-04 | 2.93E-03 |
| | multinet | 0.289 | 0.217 | 0.238 | 0.920 | 0.142 | 2.16 | 0.632 | 0.723 | 3.46E-03 | 6.98E-03 |
| cr-XVAE | corrSq | 0.302 | 0.223 | 0.246 | 0.871 | 0.133 | 2.02 | 0.549 | 0.673 | 7.17E-03 | 9.80E-03 |
| | corrAbs | 0.292 | 0.218 | 0.239 | 0.921 | 0.134 | 2.23 | 0.692 | 0.779 | 1.93E-04 | 2.95E-03 |
| | MIhist | 0.295 | 0.219 | 0.241 | 0.838 | 0.089 | 2.21 | 0.273 | 0.377 | 2.88E-01 | 3.59E-01 |
| | MIKDE | 0.294 | 0.22 | 0.241 | 0.814 | 0.115 | 2.24 | 0.513 | 0.671 | 3.66E-03 | 6.00E-03 |

**non-linear confounder**

| | | | | | | Clustering performance | | | | | |
|---|---|---|---|---|---|---|---|---|---|---|---|
| | Implementation | Reconstruction error | | | CC | Internal | | True label | | Confounder | |
| | | X1 | X2 | X1,X2 | dispersion | Silhouette | DB index | ARI | NMI | ARI | NMI |
| baseline | XVAE (unconfounded) | 0.438 | 0.278 | 0.31 | 0.957 | 0.128 | 2.17 | 0.731 | 0.821 | 1.14E-04 | 2.60E-03 |
| | KMeans | - | - | - | - | 0.160 | 1.82 | 0.251 | 0.411 | 3.25E-01 | 4.90E-01 |
| | PCA(50) + KMeans | - | - | - | - | 0.222 | 1.44 | 0.252 | 0.412 | 3.24E-01 | 4.89E-01 |
| | LR + PCA + KMeans | - | - | - | - | 0.242 | 1.48 | 0.391 | 0.567 | 2.15E-01 | 3.65E-01 |
| | XVAE | 0.227 | 0.234 | 0.232 | 0.846 | 0.123 | 2 | 0.153 | 0.293 | 4.00E-01 | 4.88E-01 |
| XVAE + FS | XVAE (corr 0.3) | 0.227 | 0.234 | 0.232 | 0.725 | 0.113 | 2.02 | 0.322 | 0.485 | 9.10E-02 | 1.16E-01 |
| | XVAE (corr 0.5) | 0.227 | 0.234 | 0.232 | 0.784 | 0.113 | 2.09 | 0.464 | 0.592 | 4.40E-02 | 5.30E-02 |
| | XVAE (pVal 0.05) | 0.227 | 0.234 | 0.232 | 0.721 | 0.192 | 1.44 | 0.226 | 0.308 | 1.16E-01 | 1.90E-01 |
| cXVAE | input | 0.224 | 0.233 | 0.23 | 0.956 | 0.146 | 1.96 | 0.477 | 0.598 | 1.95E-01 | 2.56E-01 |
| | inputEmbed | 0.217 | 0.23 | 0.225 | 0.867 | 0.145 | 2.18 | 0.758 | 0.788 | 8.64E-05 | 2.64E-03 |
| | embed | 0.216 | 0.229 | 0.224 | 0.832 | 0.114 | 2.22 | 0.498 | 0.576 | 1.34E-01 | 1.82E-01 |
| | fusedEmbed | 0.219 | 0.229 | 0.224 | 0.909 | 0.128 | 2.08 | 0.612 | 0.671 | 1.69E-01 | 2.20E-01 |
| adv-XVAE | multiclass | 0.229 | 0.235 | 0.233 | 0.951 | 0.144 | 1.930 | 0.559 | 0.637 | 1.97E-01 | 2.52E-01 |
| | multiclass (1batch) | 0.282 | 0.293 | 0.289 | 0.877 | 0.164 | 1.700 | 0.401 | 0.532 | 1.61E-01 | 3.27E-01 |
| | multiclass (scGAN) | 0.223 | 0.230 | 0.228 | 0.900 | 0.107 | 2.290 | 0.543 | 0.638 | 1.81E-01 | 2.37E-01 |
| | multinet | 0.229 | 0.245 | 0.233 | 0.951 | 0.144 | 1.930 | 0.559 | 0.637 | 1.97E-01 | 2.52E-01 |
| cr-XVAE | corrSq | 0.227 | 0.234 | 0.232 | 0.857 | 0.126 | 2.08 | 0.361 | 0.51 | 1.34E-01 | 1.93E-01 |
| | corrAbs | 0.226 | 0.234 | 0.232 | 0.818 | 0.118 | 2.03 | 0.334 | 0.496 | 1.67E-01 | 2.24E-01 |
| | MIhist | 0.228 | 0.236 | 0.233 | 0.867 | 0.126 | 2.01 | 0.336 | 0.493 | 1.91E-01 | 2.59E-01 |
| | MIKDE | 0.227 | 0.234 | 0.231 | 0.852 | 0.117 | 2.10 | 0.337 | 0.487 | 1.92E-01 | 2.53E-01 |

Supplementary Table 2: **Performance overview of different implementations of each deconfounding strategy for single confounders -** continued

**categorical confounder**

| | Implementation | Reconstruction error | | | CC | Clustering performance | | | | | |
|---|---|---|---|---|---|---|---|---|---|---|---|
| | | | | | | Internal | | True label | | Confounder | |
| | | X1 | X2 | X1,X2 | dispersion | Silhouette | DB index | ARI | NMI | ARI | NMI |
| baseline | XVAE (unconfounded) | 0.438 | 0.278 | 0.31 | 0.957 | 0.128 | 2.17 | 0.731 | 0.821 | 1.14E-04 | 2.60E-03 |
| | KMeans | - | - | - | - | 0.150 | 1.91 | 0.0610 | 0.117 | 2.18E-01 | 3.71E-01 |
| | PCA(50) + KMeans | - | - | - | - | 0.197 | 1.62 | 0.0631 | 0.120 | 2.15E-01 | 3.68E-01 |
| | LR + PCA + KMeans | - | - | - | - | 0.186 | 1.71 | 0.150 | 0.269 | 7.09E-02 | 1.52E-01 |
| | XVAE | 0.203 | 0.225 | 0.217 | 0.755 | 0.0819 | 2.32 | 0.335 | 0.476 | 3.31E-03 | 6.50E-03 |
| XVAE + FS | XVAE (corr 0.3) | 0.203 | 0.225 | 0.217 | 0.727 | 0.117 | 1.98 | 0.212 | 0.307 | 7.05E-03 | 1.23E-02 |
| | XVAE (corr 0.5) | 0.203 | 0.225 | 0.217 | 0.761 | 0.0842 | 2.31 | 0.339 | 0.482 | 4.00E-03 | 7.38E-03 |
| | XVAE (pVal 0.05) | - | - | - | - | - | - | - | - | - | - |
| cXVAE | input | 0.197 | 0.217 | 0.209 | 0.66 | 0.106 | 2.33 | 0.0961 | 0.129 | 2.36E-01 | 4.14E-01 |
| | inputEmbed | 0.195 | 0.218 | 0.209 | 0.873 | 0.0877 | 2.43 | 0.443 | 0.53 | 1.47E-03 | 5.22E-03 |
| | embed | 0.195 | 0.216 | 0.208 | 0.9 | 0.109 | 2.43 | 0.657 | 0.742 | 8.35E-04 | 3.65E-03 |
| | fusedEmbed | 0.191 | 0.216 | 0.206 | 0.829 | 0.101 | 2.38 | 0.575 | 0.680 | 3.76E-04 | 2.96E-03 |
| adv-XVAE | multiclass | 0.204 | 0.225 | 0.217 | 0.707 | 0.123 | 2.15 | 0.254 | 0.298 | 1.29E-01 | 2.48E-01 |
| | multiclass (1batch) | 0.263 | 0.286 | 0.277 | 0.777 | 0.146 | 2.05 | 0.140 | 0.193 | 1.36E-01 | 3.37E-01 |
| | multiclass (scGAN) | 0.196 | 0.218 | 0.210 | 0.681 | 0.111 | 2.41 | 0.0115 | 0.0102 | 3.58E-01 | 5.92E-01 |
| | multinet | - | - | - | - | - | - | - | - | - | - |
| cr-XVAE | corrSq | 0.200 | 0.222 | 0.213 | 0.744 | 0.079 | 2.380 | 0.206 | 0.314 | 3.78E-02 | 9.69E-02 |
| | corrAbs | 0.201 | 0.223 | 0.214 | 0.704 | 0.093 | 2.410 | 0.205 | 0.279 | 1.31E-01 | 2.51E-01 |
| | MIhist | 0.204 | 0.223 | 0.215 | 0.755 | 0.090 | 2.310 | 0.393 | 0.532 | 2.09E-03 | 6.24E-03 |
| | MIKDE | 0.202 | 0.225 | 0.216 | 0.815 | 0.091 | 2.280 | 0.319 | 0.469 | 2.15E-03 | 5.81E-03 |

**multiple confounders**

| | | reconstruction error | | | CC | Clustering performance | | | | | | | | | |
|---|---|---|---|---|---|---|---|---|---|---|---|---|---|---|---|
| | | | | | | Internal | | True label | | Linear conf. | | Non-linear conf. | | Categorical conf. | |
| | | X1 | X2 | X1,X2 | dispersion | Silhouette | DB index | ARI | NMI | ARI | NMI | ARI | NMI | ARI | NMI |
| XVAE | mean | 0.145 | 0.174 | 0.161 | 0.725 | 0.077 | 2.441 | 0.216 | 0.269 | 1.50E-02 | 2.57E-02 | 1.40E-01 | 1.97E-01 | 6.72E-02 | 1.33E-01 |
| | std | 0.004 | 0.002 | 0.003 | 0.043 | 0.006 | 0.058 | 0.089 | 0.100 | 1.93E-02 | 3.03E-02 | 4.33E-02 | 4.71E-02 | 4.77E-02 | 8.20E-02 |
| XVAE + FS | mean | 0.145 | 0.174 | 0.161 | 0.731 | 0.089 | 2.341 | 0.265 | 0.327 | 7.17E-03 | 1.20E-02 | 1.91E-02 | 2.78E-02 | 1.09E-01 | 2.03E-01 |
| | std | 0.004 | 0.002 | 0.003 | 0.037 | 0.006 | 0.064 | 0.085 | 0.104 | 8.63E-03 | 1.06E-02 | 2.99E-02 | 3.97E-02 | 5.75E-02 | 9.60E-02 |
| cXVAE | mean | 0.130 | 0.159 | 0.146 | 0.905 | 0.126 | 2.243 | 0.634 | 0.713 | 1.25E-04 | 2.69E-03 | -3.67E-04 | 2.18E-03 | 4.62E-04 | 3.35E-03 |
| | std | 0.003 | 0.002 | 0.002 | 0.022 | 0.009 | 0.065 | 0.042 | 0.031 | 3.83E-04 | 5.10E-04 | 4.68E-04 | 5.78E-04 | 5.01E-04 | 7.26E-04 |
| adv-XVAE | mean | 0.141 | 0.172 | 0.158 | 0.753 | 0.106 | 2.207 | 0.225 | 0.262 | 1.56E-02 | 2.58E-02 | 1.07E-01 | 1.51E-01 | 1.06E-01 | 2.22E-01 |
| | std | 0.003 | 0.005 | 0.004 | 0.066 | 0.008 | 0.054 | 0.120 | 0.130 | 2.28E-02 | 3.48E-02 | 5.23E-02 | 7.37E-02 | 5.14E-02 | 1.04E-01 |
| cr-XVAE | mean | 0.145 | 0.174 | 0.161 | 0.764 | 0.095 | 2.234 | 0.369 | 0.490 | 2.67E-03 | 6.91E-03 | 6.88E-03 | 1.71E-02 | 4.87E-04 | 3.47E-03 |
| | std | 0.005 | 0.002 | 0.003 | 0.031 | 0.007 | 0.074 | 0.064 | 0.064 | 1.74E-03 | 3.58E-03 | 1.05E-02 | 2.44E-02 | 5.02E-04 | 7.75E-04 |

Supplementary Table 3: **Overview performances of deconfounding strategy for multiple confounder simulations**
